# SARS-CoV-2 variants from long-term, persistently infected immunocompromised patients have altered syncytia formation, temperature-dependent replication, and serum neutralizing antibody escape

**DOI:** 10.1101/2024.05.18.594818

**Authors:** Camille Wouters, Jaiprasath Sachithanandham, Elgin Akin, Lisa Pieterse, Amary Fall, Thao T. Truong, Jennifer Dien Bard, Rebecca Yee, David J. Sullivan, Heba H. Mostafa, Andrew Pekosz

## Abstract

SARS-CoV-2 infection of immunocompromised individuals often leads to prolonged detection of viral RNA and infectious virus in nasal specimens, presumably due to the lack of induction of an appropriate adaptive immune response. Mutations identified in virus sequences obtained from persistently infected patients bear signatures of immune evasion and have some overlap with sequences present in variants of concern. We characterized virus isolates from two COVID-19 patients undergoing immunosuppressive cancer therapy, with all isolates obtained greater than 100 days after the initial COVID-19 diagnoses and compared to an isolate from the start of the infection. Isolates from an individual who never mounted an antibody response specific to SARS-CoV-2 despite the administration of convalescent plasma showed slight reductions in plaque size and some showed temperature-dependent replication attenuation on human nasal epithelial cell culture compared to the virus that initiated infection. An isolate from another patient - who did mount a SARS-CoV-2 IgM response – showed temperature dependent changes in plaque size as well as increased syncytia formation and escape from serum neutralizing antibody. Our results indicate that not all virus isolates from immunocompromised COVID-19 patients display clear signs of phenotypic change, but increased attention should be paid to monitoring virus evolution in this patient population.

## Introduction

The evolution of novel SARS-CoV-2 antigenic variants has reduced the effectiveness of current vaccines and monoclonal antibody treatments, contributing to sustained SARS-CoV-2 transmission [1], [2]. SARS-CoV-2 has a relatively low genome mutation rate compared to RNA viruses such as influenza and HIV, due to a proofreading exoribonuclease encoded by coronaviruses [3]. This in combination with narrow transmission bottlenecks means very little genetic diversity is generated and transmitted on to new hosts during typical acute infections [4], [5]. However, during prolonged infections in immunocompromised patients (ICPs), the appearance and disappearance of mutations is observed within days to weeks, and is often associated with the presence of infectious virus at late times post infection [1], [6], [7], [8], [9], [10], [11], [12], [13]. These infections are distinct from infections after which SARS-CoV-2 RNA positivity continues in the absence of infectious virus, and with no significant virus genome mutations [9], [11], [14]. Persistently low levels of Spike antibodies in ICPs could promote the selection of new virus variants over the course of continued replication cycles within the host [15]. ICPs often develop reduced antibody responses after SARS-CoV-2 infection or vaccination [9], [13], [16], [17], [18], [19], [20]. Rapid changes in variant composition within an individual suggest the selection for variants containing certain mutations that promote increased replication fitness, escape from anti-SARS-CoV-2 antibodies or plasma administered therapeutically, or both [3]. In support of this observation, monoclonal antibody or convalescent plasma therapy in ICPs has corresponded to increased frequencies of mutations in the Spike protein [1], [3], [6], [7], [9], [20], and SARS-CoV-2 evolving in an immunocompromised HIV patient was only weakly neutralized by the patient’s own plasma [21].

Mutations in variants isolated over the course of persistent infections are reflected in global variants of concern, and Alpha and Omicron variants have been hypothesized to have evolved in immunocompromised persons [3], [22], [23], [24]. While most variants emerging in immunocompromised individuals do not appear to be transmitted, the direct forward transmission of an Omicron BA.1 sub-lineage which acquired 8 additional Spike mutations in an ICP to three other ICPs and 2 immunocompetent individuals has been reported [25]. Variants appearing in ICPs have not been carefully studied for their replication and escape from pre-existing immunity, which is essential to gauge the potential risk of these variants to the general population. While sequence analysis may predict some phenotypic changes such as escape from neutralizing antibodies, it cannot predict the overall replication fitness of the emerging variants – that assessment requires characterization of patient-derived virus isolates. Replication fitness comparisons among isolates from persistently infected ICPs are limited and have so far only involved immortalized cell lines at a single temperature [26], [27]. Understanding how SARS-CoV-2 populations change within an immunocompromised host informs us of viral and host factors driving selection at the origin of potential new variants.

We isolated and sequenced SARS-CoV-2 from infections in three immunocompromised B cell acute lymphocytic leukaemia patients between May and November 2020 [9]. This initial study indicated that Patient 1 did not have culturable virus by two weeks after symptom onset and is therefore excluded from this paper [9]. Patient 2’s Day 0 virus was collected from a nasal swab obtained before symptoms began, but after exposure to a SARS-CoV-2-positive contact [9]. Patient 2 received CD19-directed CAR-T cell therapy prior to their infection, had a CD4/CD8 ratio <1 (associated with altered immune function) and had no detectable antibodies against SARS-CoV-2 until the regular approximately weekly administration of convalescent plasma therapy starting from day 103 onwards (plasma was also administered once at day 78) [9], [28]. Patient 3 Day 0 virus was collected soon after fever onset. Patient 3 was receiving chemotherapy, had CD4/CD8 ratio <1, and from day 80 post-infection had detectable IgM antibodies to SARS-COV-2 Spike, with no evidence of a switch to IgG [9].

To understand the impact of SARS-CoV-2 mutations appearing during prolonged infection of ICPs, we characterized virus isolates for changes in temperature-dependent replication in transformed and primary cell cultures, syncytia formation and escape from serum neutralizing antibodies. In this way, we could determine the overall changes in virus phenotypes that resulted from an accumulation of mutations across the viral genome in addition to measuring specific changes in Spike protein function and neutralizing antibody escape.

## Methods

### Institutional Review Board Approvals

For convalescent plasma, donor specimens were obtained with written informed consent per the protocols approved by the institutional review boards at Johns Hopkins University School of Medicine (IRB00248402 donor and IRB00247590 early treatment) as single Institutional Review Board for all participating sites and the Department of Defense Human Research Protection Office. Virus isolation was performed on deidentified samples under Johns Hopkins protocol number IRB00288258.

### Cell Culture

VeroE6-Transmembrane Serine Protease 2 (TMPRSS2) overexpressing cells (Vero/TMPRSS2) (cell repository of the National Institute of Infectious Diseases, Japan) [29], Vero E6-TMPRSS2-T2A-ACE2 cells overexpressing ACE2 (Vero/TMPRSS2/ACE2) (BEI Resources, NIAID, NIH) and Lenti-X HEK 293T cells (TakaraBio) were cultured at 37°C and 5% CO2 in complete cell culture media (CM; DMEM supplemented with 1% GlutaMAX (Life Technologies, Cat#35050061), 10% Fetal Bovine Serum (FBS; Gibco, Cat#26140079), 1% Penicillin/Streptomycin mixture (Quality Biologicals, Cat#381 120-095-721), and 1% 100mM sodium pyruvate solution (Sigma, Cat#S8636-100ML)). Human nasal epithelial cells (HNEpC; PromoCell, Cat#C-12620) were expanded to confluency with PneumaCultTM Ex Plus Media (StemCell, Cat#05040) at 37°C and 5% CO2 on Transwell insert (Corning Cat#3470). Confluent cells were fully differentiated in ALI (air-liquid interface) with PneumaCult ALI Basal Medium (Stemcell, Cat#05002) and 1X PneumaCult ALI Supplement (Stemcell, Cat#05003). 1% PneumaCult ALI Maintenance Supplement (Stemcell, Cat#05006), 0.5% Hydrocortisone stock solution (Stemcell, Cat#07925) and 0.2% Heparin solution (Stemcell, Cat#07980) were added to the ALI Basal Medium.

### Virus Plaque Picking, Seed Stock and Working Stock Generation

All work with live SARS-CoV-2 virus was performed under Biosafety Level 3 (BSL-3) conditions using Institution approved procedures. Virus isolates derived from nasal swabs [9] were serially diluted 10-fold, and 6-well Vero/TMPRSS2 plates were infected with virus dilutions. After a 1-hour incubation at 37°C, a 1% agarose/1x Modified Eagle Medium (MEM, Gibco) overlay was added. After approximately 4 days, distinct virus plaques were picked using a P1000 pipette tip and resuspended in 500 μL IM. 150 μL of this suspension was used to inoculate a single well of a 24-well plate containing 350 μL IM. Cells were monitored daily for cytopathic effect (CPE) and supernatants were harvested when CPE was visible and > 75% of cells were detached. 140 μL supernatant was inactivated using Triton X-100 to a final concentration of 0.5% for downstream RNA extraction and sequencing, and the remaining supernatant was frozen as plaque purified seed stock.[11]

The Spike sequences of seed stocks were determined to choose plaques for downstream working stock generation and virus characterisation. RNA was extracted suing the QIAamp 96 Viral RNA Kit. Spike PCR was carried out using Super Script III One-Step RT-PCR System with Platinum Taq High Fidelity DNA Polymerase (Thermo Fisher Scientific, Waltham, USA) with Spike forward (F1) and reverse (R2) primers (see below table for sequences). The amplified PCR products were purified using the QIAquick PCR Purification Kit by following the manufacturer’s instructions and submitted to JHMI Synthesis & Sequencing Facility for Sanger sequencing using the following 7 Spike forward and reverse primers (Supplementary table 1).

Seed stocks containing Spike mutations most closely resembling the majority SNPs from the origin patient nasal swab RNA results were used to grow up working stocks of virus. These stocks were then used for amplicon based whole viral genome sequencing to establish the consensus sequence and frequency of SNPs in the working stock (see below methods) (Fig. 1). To generate a working stock, 80% confluent flasks of Vero/TMPRSS2 cells were infected at 33°C at an MOI of 0.05 in 7 ml IM. After 1 hour, an additional 10 ml IM was added to the flasks. The flasks were incubated until 75% CPE was observed. Supernatant was collected and centrifuged at 800 x*g* for 5 minutes to remove cell debris. The supernatant was then aliquoted and stored at -65°C as working stock (henceforth referred to as an isolate) [30], [31].

**Figure 1:**
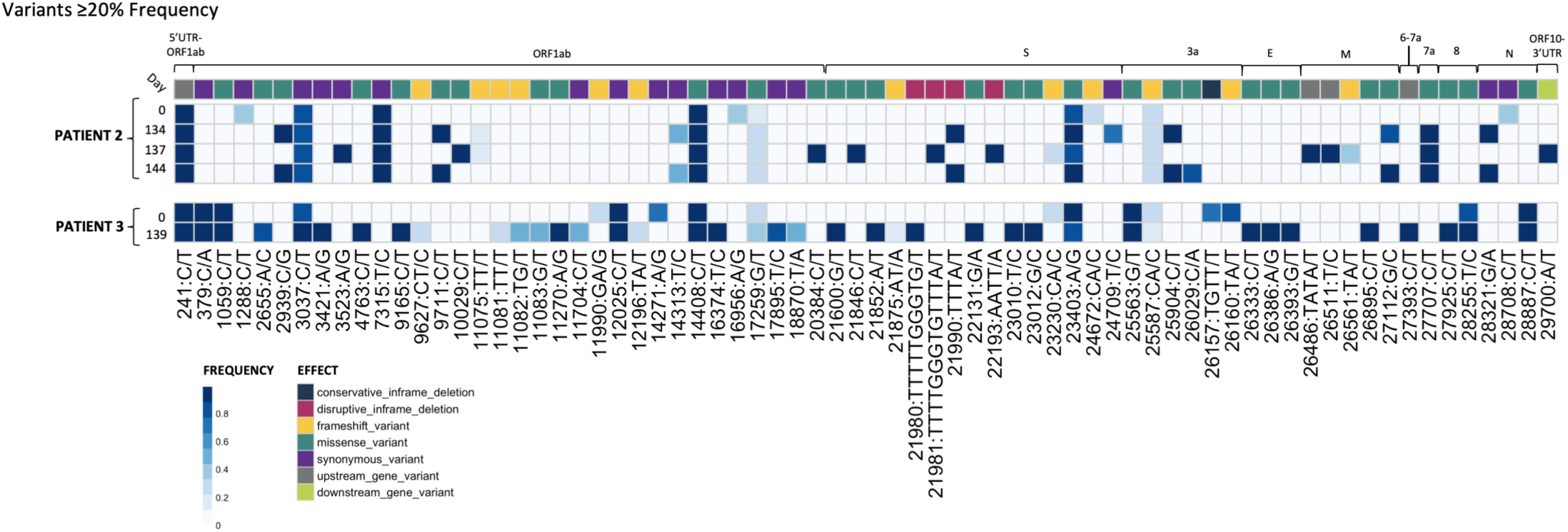
Plaque picked isolates from Patients 2 and 3 show the accumulation of mutations during prolonged SARS-CoV-2 replication in the patients. RNA-sequencing results from plaque-picked SARS-CoV-2 virus working stocks used for characterisation work in this paper (UTR, untranslated region; S, spike; E, envelope; M, matrix; N, nucleocapsid). All mutations are relative to the 2020 Wuhan-Hu-1 reference genome (NCBI Reference Sequence: NC_045512.2). For comparison to nasal swab sequences from [9], see Supplementary Tables 1 and 2.

### Sequencing of Plaque Picked SARS-CoV-2 Isolates

Viral RNA was extracted, sequenced and variants were called as previously described [32], [33]. Briefly, variants were called using the arctic-ncov2019 medaka protocol against reference hCoV-19/Wuhan/WIV04/2019 (EPI_ISL_402124). Variants were manually inspected against BAM files using Integrated Genomics Viewer (v2.12.3) and Geneious Prime (2023.1.2 Build 2023-04-27). Resulting variant call files (VCFs) were indexed and merged using tabix (v1.17) and bcftools (v1.17). Merged VCFs were filtered for quality (QUAL ≥ 30) and mono-allelic variant calls. Allele frequency was calculated as the abundance of alternate allele reads over reference allele reads using vcf2pmatrix.py and ratio.py. A bi-allelic tandem repeat insertion variant at position 11,074 CT/CCT,C was removed due to visualization constraints and can be viewed in the: Merged_PASS_complete_calls.vcf. Variants were visualized using custom scripts and the pheatmap package (v1.0.12). All scripts are available at https://github.com/Pekosz-Lab/Wouters_2024_CHLA. Variant calls at a frequency below 0.2 were excluded from Fig. 1. All mutations are relative to the 2020 Wuhan-Hu-1 reference genome (NCBI Reference Sequence: NC_045512.2).

### Tissue Culture Infectious Dose (TCID50) Assay

SARS-CoV-2 infectious virus titers were determined by TCID_50_ [30], [31]. Vero/TMPRSS2 cells were grown on 96-well plates until 80% confluence. Cells were washed with 1x PBS supplemented with 0.1 grams/litre CaCl_2_ and MgCl_2_, and 180 μL IM was added to each well. Virus samples were serially diluted 10-fold, and each diluted sample was added in sextuplicate to the 96-well plates. The plates were incubated for 5 days at 37°C and then fixed in 2% formaldehyde, followed by staining with Napthol Blue Black. TCID_50_ values were calculated using the Reed-Meunch method [34].

### Vero/TMPRSS2 Infections

Vero/TMPRSS2 cells were grown on 24-well plates to 100% confluency, washed once with IM and infected at an MOI of 0.01 [30], [31]. Four replicate wells were infected per virus. Plates were incubated at 33°C or 37°C for 1 hour, washed with IM, and 500 μL was replaced onto the cells. At the indicated times post infection, supernatants were collected and stored at -65°C for TCID50 determination, and fresh IM was added.

### Human Nasal Epithelial Cell (hNEC) Infections

The apical side of the hNEC Transwell was washed three times with 1x PBS with a 10-minute incubation at 37°C during each wash step [30], [31]. Diluted virus was added to the apical side at an MOI of 0.05 in 100 μL IM. After a 2 hour incubation at 33°C or 37°C, the apical side was washed three times with 1x PBS. At every timepoint post infection, 100 μL IM was added to the apical side, incubated for 10 minutes at 33°C or 37°C and harvested as supernatant for TCID_50_ determination. Basolateral media was replaced every 48 hours. Four wells were used per virus per independent hNEC experiment. Occasionally, hNEC wells were not infected after incubation with virus at an MOI of 0.05. In these instances, uninfected wells were excluded from the growth curve data.

### Plaque Reduction Neutralisation Test (PRNT)

Donor convalescent plasma samples collected between July-November 2020 with known NT50 values against ancestral Washington-1 (SARS-CoV-2/USA-WA1/2020), Delta (hCoV19/USA/MD-HP05660/2021), and Omicron (hCoV19/USA/MD-HP20874/2021) variants were selected for PRNTs using the isolates from patients 2 and 3 [35]. Convalescent plasma samples were heat inactivated by incubation at 56°C for 1 hour. PRNTs were then run at 37°C as previously described [30], [36]. GraphPad Prism 9 was used generate inhibition dose-response curves from plaque forming unit counts, and IC_50_ values were calculated using a non-linear regression model.

### Spike Plasmid Preparation

SARS-CoV-2 virus stocks were inactivated by incubation in a final concentration of 0.5% NP-40 for 30 minutes. RNA was extracted using QIAamp Viral RNA Mini Kit (Qiagen), and cDNA was produced using a ProtoScript® II Reverse Transcriptase (New England Biolabs) reaction and a Spike-specific reverse primer (5’ CTGAAGGAGTAGCATCCTTG 3’). The SARS-CoV-2 Spike coding region was then amplified using Q Hot Start High-Fidelity DNA Polymerase (New England Biolabs) with forward (5’ TCATCGATGCATGGTACGCCACCATGTTTGTTTTTCTTGTTTTATTG 3’) and reverse (5’ CTGCTAGCTCGAGCATGTTATGTGTAATGTAATTTGACTCC 3’) primers. The product of this reaction was then run on a 0.8% agarose gel and purified using Zymoclean Gel DNA Recovery Kit (Zymo Research) to yield the final Spike DNA fragment. Empty pCAGGS plasmid vector was digested using restriction enzymes KpnI-HF and SphI-HF (New England Biolabs) and purified using QIAquick PCR Purification Kit (Qiagen) [37]. Spike DNA fragments were introduced into the digested pCAGGS vector using NEBuilder® HiFi DNA Assembly (New England Biolabs). The product of this assembly reaction was transformed into 5-alpha Competent *E. coli* (New England Biolabs), which were plated and incubated at 37°C overnight on LB Agar Carbenicillin (100 µg/mL) plates. Picked colonies were grown up overnight in LB-Carbenicillin (100 µg/mL). Whole plasmids from single colonies were sequenced to confirm that the Spike sequence within the pCAGGS plasmids was identical to the most common SNPs contained within plaque purified virus isolates. Five Spike-pCAGGS plasmids were generated for the six total Patient 2 and 3 isolate plasmids characterised, as Patient 2 Day 0 and Patient 3 Day 0 Spike proteins have identical sequences.

### Flow Cytometry for Surface Spike

Vero/TMPRSS2 cells were plated for 90% confluency in 6-well plates 24 hour before Spike transfection. Immediately before transfection, CM was replaced with Opti-MEM reduced serum media (Gibco). Each well was transfected with 2.5 μg Spike-pCAGGS plasmid using TransIT®-LT1 Transfection Reagent (Mirus). 24 hours after transfection, the Opti-MEM was removed and cells were trypsinised in 500 μL 1x 0.5% Trypsin-EDTA (Life Technologies). 500 μL CM was then added, and the cells were pelleted at 200 x g for 4 minutes (all washes prior to cell fixation were conducted using these centrifuge settings). The cells were washed three times with 1X PBS and resuspended in PBS. Dead cells were then stained using the Live/Dead Fixable Aqua Dead Cell Stain Kit (Thermofisher). After a 30-minute incubation, cells were washed once in 1x PBS and once in Flow Buffer (1% BSA in 1x PBS) (BSA from Sigma-Alrich). Cells were incubated for 20 minutes at room temperature in the primary antibody SARS-CoV-2 (2019-nCoV) Spike S2 Antibody Chimeric MAb (Sinobiological RRID Number: AB_2857932), diluted 1:75 in Flow Buffer. Cells were then washed once in Flow Buffer, followed by secondary antibody staining. Cells were incubated for 20 minutes at room temperature in Goat anti-Human IgG (H+L) Cross-Adsorbed Secondary Antibody, Alexa Fluor™ 647 (Invitrogen) diluted 1:1000 in Flow Buffer to 2 μg/ml. Cells were washed once more with Flow Buffer and once more with 1X PBS before fixation in 4% paraformaldehyde for 30 minutes. After fixation, all washes were conducted at 500 x g for 4 minutes. Cells were washed twice with Flow Buffer, and then resuspended in Flow Buffer. Samples were run on a BD LSRII machine, and flow cytometry gating was conducted using FlowJo 10. Cells positive for surface Spike were gated from live single cells (Supplementary Figure 1a).

#### mCherry Lentivirus Production

The pLV lentivirus transfer plasmid (VectorBuilder) backbone (containing Blasticidin resistance gene for the selection of transduced cells) and mCherry gene were PCR amplified using Q5® Hot Start High-Fidelity DNA Polymerase (New England Biolabs), remnant parent templates were digested using DpnI (New England Biolabs), and DNA products were gel purified using Zymo Gel DNA Recovery Kit (Zymogen). The mCherry gene was cloned into the pLV plasmid using using NEBuilder® HiFi DNA Assembly (New England Biolabs), to generate the final pLV-mCherry product. The product of this assembly reaction was transformed into 5-alpha Competent *E. coli* (New England Biolabs), which were plated and incubated at 37°C overnight on LB Agar Carbenicillin (100 µg/mL) plates. Picked colonies were grown overnight in LB-Carbenicillin (100 µg/mL), and the final pLV-mCherry plasmid was confirmed by whole plasmid sequencing.

Lenti-X HEK 293T cells were plated in 6 well plates for 90% confluency. Each well was transfected with a mixture of the following: 150 ul jetPRIME buffer (Polyplus), 6 ul jetPRIME transfection reagent (Polyplus), 0.25 μg psPAX2 packaging plasmid (AddGene #12260), 0.25 μg pCMV-VSV-G envelope plasmid (AddGene #8454), 1 μg mCherry-pLV. 1 day after transfection, media was replaced with fresh CM. 3 days after transfection, cell supernatant containing lentivirus was collected and centrifuged at 500 x g for 5 minutes to pellet cell debris. The mCherry lentivirus was stored at -65°C until use.

#### Lentivirus Transduction and Clonal Cell Selection for Stable Expression of mCherry in Vero/TMPRSS2 Cells

Vero/TMPRSS2 cells were plated at 50% confluency in 6-well plates. 24 hours after plating, media was removed from the cells and replaced with 1 ml of mCherry lentivirus supernatant mixed with 8 ug Polybrene Infection / Transfection Reagent (Sigma-Aldrich). 24 hours after lentivirus addition, media was replaced with fresh CM. 3 days post lentivirus addition, CM was replaced with CM containing 2 ug/ml Blasticidin to select for successfully transduced cells. Cells were transferred to T75 flasks after reaching confluency in 6-well plates, and 2 weeks after blasticidin addition, were plated onto 100 mm petri dishes at low density to enable clonal cell isolation using cloning cylinders. The clone with the brightest mCherry expression was expanded for use in syncytia assays, and is now termed Vero/TMPRSS2/mCherry. Vero/TMPRSS2/mCherry cells were maintained in CM containing 2 ug/ml Blasticidin.

### Syncytia Assay

Vero/TMPRSS2/mCherry cells were plated at 90% confluency. 24 hours after plating, media was changed to OptiMEM. The Vero/TMPRSS2/mCherry cells were transfected with pCAGGS-Spike plasmids using TransIT®-LT1 Transfection Reagent (Mirus). Vero/TMPRSS2/ACE2 cells were incubated with 10 μM CellTracker Green CMFDA Dye (Invitrogen) for 30 minutes at room temperature. 5 hours after pCAGGS-Spike transfection, the transfected Vero/TMPRSS2/mCherry cells were mixed at a 1:1 ratio with the CMFDA-treated Vero/TMPRSS2/ACE2 cells, and plated onto 8-well chamber slides (Ibidi) at a total density of 7 x 10^4^ cells/cm^2^. 24 hours after plating, the slides were washed three times in 1x PBS, fixed in 4% paraformaldehyde, and washed once in 1x PBS. Nuclei were stained for 5 minutes in 5 μg/ml Hoechst 33258 dye (Thermofisher), and washed twice more in 1x PBS. Wells were imaged in 1x PBS.

The entire area of each well was imaged in tiles using a Leica Thunder imaging system at 10x magnification. Raw images of blue (nuclei), red and green channels were then used for analysis in CellProfiler. A custom CellProfiler pipeline was used to determine the number of nuclei contained within syncytia, defined as areas with red and green fluorescence. Pipeline settings are available in the raw data folder for the syncytia assay. Images containing well edges were excluded from downstream analysis as there was significant overlap in red and green channels in that area of the slide. The average size of nuclei was determined by dividing the total nuclei area by the total number nuclei in un-transfected control wells, as syncytia containing overlapping nuclei decreased the accuracy of nuclei counts in Spike-transfected wells. For the same reason, the percentage of nuclei in syncytia was calculated using total nuclei counts from mock wells. Flow cytometry revealed no statistically significant differences in Spike expression between the pCAGGS-Spike plasmids, and therefore syncytia assay results were not normalised to Spike expression data.

### Statistical Analyses

All statistical analysis was performed using GraphPad Prism 10. PRNT data was assumed to be normally distributed and was matched by serum sample. Syncytia assay and flow cytometry data was assumed to be normally distributed, and individual experimental repeats were treated as matched sets to account for experiment-to-experiment variability.

## Results

### Plaque picked virus stocks have multiple mutations compared to the initial infecting virus that align with global Variants of Concern

Four Patient 2 nasal swabs (Day 0, 134, 137 and 144) and two Patient 3 nasal swabs (Day 0 and 139) were chosen for characterisation, as these sequences displayed multiple genetic changes across the genome when compared to the initial infecting virus [9]. All Patient 2 specimens matched to Nextstrain clade 20A, and all Patient 3 specimens matched to clade 20C [9]. Day 0 isolate from each patient served as a parental reference for every experiment to represent the virus at the start of the persistent infection [11]. Virus isolated from nasal swab samples was used to generate plaque purified seed stocks of the patient viruses. The working stocks generated from the plaque purified seed stocks were sequenced to assess differences in SNP frequency across the entire SARS-CoV-2 genome between the plaque picked isolate versus the infecting virus ([9]Fig. 1 and Supplementary Tables 2 and 3).

Seventy mutations at different sites within the SARS-CoV-2 genome (as compared to the Wuhan-Hu-1 reference) were found at a frequency of 0.2 or higher in plaque picked isolate working stocks (Fig. 1). 20 of these mutations were not detected in the original patient nasal swab samples [9], and 6 of these were found at an allele frequency > 0.5. 12 of these 20 unexpected mutations were frameshift mutations, though only 2 were present at >0.5 frequency. All mutations identified in the corresponding nasal swabs at frequencies > 0.5 were present in the Patient 3 Day 0 and 139 and Patient 2 Day 0, 134 and 137 plaque picked isolates. However, in the Patient 2 Day 144 plaque picked isolate working stock, 6 mutations ranging from 47% - 59% frequency within the nasal swab samples were lost, indicating that the Day 144 isolate represents one sequence from a mixed population that existed in Patient 2 at day 144. As a result, only two mutations were found to distinguish Patient 2 Day 134 and Day 144 isolate sequences from each other, at sites 24709 (Spike protein, synonymous mutation) in Day 134 and ORF3a substitution Q213K (26029 C/A) in Day 144 (Fig. 1), with neither mutation detected in the nasal swab sequence [9]. Other mutations present in the nasal swab that distinguished Day 134 and Day 144 viruses from each other in the nasal swab were lost during plaque picking and working stock generation, notably including a non-synonymous Spike mutation at 21990 (Spike T22I) which was lost in all 4 sequenced plaques picked before working stock generation and whole genome RNA-seq [9]. The majority (73%) of Spike mutations found in the isolates were at > 85% frequency within the virus stock (Fig. 1).

Some mutations appearing in nasal swab viruses and plaque picked isolates at later infection timepoints are identical to ones appearing months to years later in global SARS-CoV-2 variants including Alpha, Delta and Omicron. Spike mutations appearing in plaque picked isolates from Patient 2 Day 134, 137 and 144 swabs and the Patient 3 Day 139 swab include changes at amino acids L141-Y145 (ΔL141-V143 (21980 TTTTTGGTG/T), ΔL141-Y144 (21981 TTTTGGGTGTTTA/T) and ΔY145 (TTTA/T)) mutated or deleted in Alpha and Omicron, and E484 (23012 G/C) mutated in Beta and Omicron variants [9], [38], [39]. Significantly, deletions at ΔL141-144 have also been recorded in at least 6 separate case studies of persistently infected ICPs [9], [40].

Mutations in non-Spike ORFs within the 30 kB SARS-CoV-2 genome can impact viral fitness [41], [42], [43], [44]. For example, the commonly occurring ORF7a C-terminal truncation attenuates virus-mediated interferon response suppression [42]. Non-Spike mutations that appear in the nasal swabs and plaque picked isolates include an ORF7a A105V (27707 C/T) mutation (which appears and persists in all late time point Patient 2 viruses), ORF8 T11I (27925 C/T, in Patient 3 Day 139), and nsp4 T3255I (10029 C/T in Patient 2 Day 137 virus) (Fig. 1) [9]. This nsp4 T3255I (10029 C/T) mutation appeared for Patient 2 Day 137 virus months before it became dominant in SARS-CoV-2 GISAID sequences and has been found in all global variants since mid-2021 [38], [45]. Likewise, ORF8 T11I (28255 C/T) briefly peaked at 15% of United States GISAID sequences and was a defining mutation of the Iota lineage [38], [45]. The alignments between the virus non-Spike mutations and those in widespread SARS-CoV-2 variants suggest these mutations may confer some competitive advantage to the virus within a persistently infected host.

### Patient 2 virus isolates have distinct plaque sizes and temperature dependent replication differences on Vero/TMPRSS2 cells and hNECs

Vero/TMPRSS2 cells are highly permissive to SARS-CoV-2 replication and are widely used to investigate differences in SARS-CoV-2 growth kinetics and plaque formation between variants [29], [46]. SARS-CoV-2 replication kinetics vary according to temperature, and Vero/TMPRSS2 growth curves and plaque assays were conducted at 33°C and 37°C to represent the range of temperatures within the human respiratory tract [47], [48]. Patient 2 Day 137 and Day 144 isolates have smaller plaque sizes on Vero/TMPRSS2 cells when compared to the Day 0 virus at 33°C and 37°C (Fig. 2 a-d). The Day 134 isolate, despite being isolated from a swab taken only days earlier than Day 137, showed no differences in plaque size versus Day 0 virus at 33°C. However, Day 134 plaques were visibly smaller than Day 0 plaques at 37°C. Overall, there was a trend of decreasing plaque size in later timepoint viruses versus Day 0 isolate.

**Figure 2:**
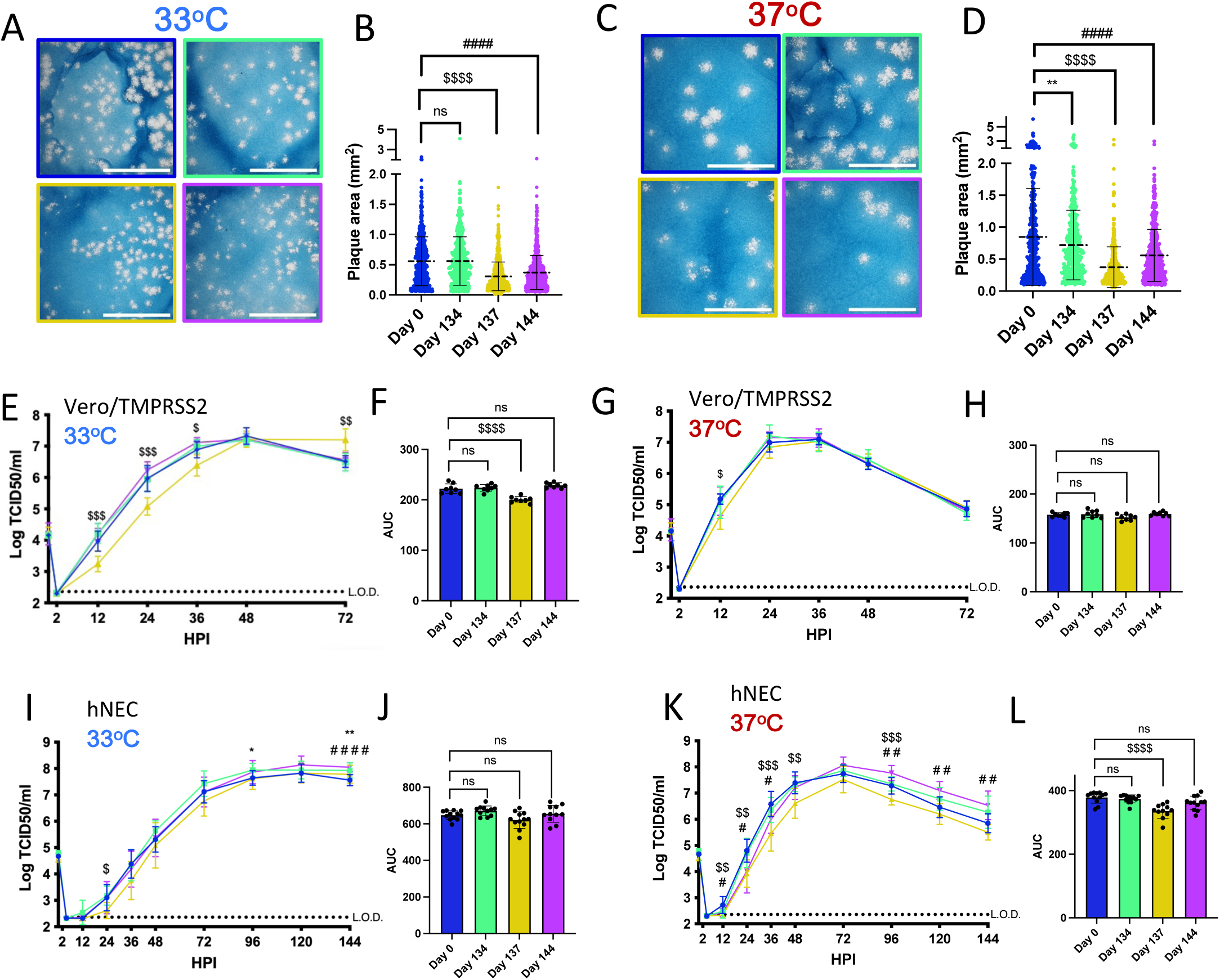
Viruses isolated from Patient 2 later during the infection have distinct temperature-dependent phenotypes compared to Day 0 virus. Comparisons: * (Day 0 to Day 134), $ (Day 0 to Day 137), # (Day 0 to Day 144). p values displayed as: “ns” p > 0.05,), * p < 0.05, ** p < 0.01, *** p < 0.001, ****p <0.0001. A, C: Representative images of plaques from each virus isolate at 33°C and 37°C respectively, in Vero/TMPRSS2 cells. Scale bar = 10 mm. B, D: Quantified plaque sizes, >500 (33°C) and >390 (37°C) plaques per virus accumulated from 3 independent experiments, one-way ordinary ANOVA with Bonferroni’s multiple comparisons test (all comparisons to Day 0 isolate). E, G: Growth curves showing virus replication on Vero/TMPRSS2 cells at 33°C and 37°C respectively, the data are derived from 2 independent experiments with four wells per virus per experiment, standard deviation shown on error bars, two-way repeated measures ANOVA with Bonferroni’s multiple comparisons test (all comparisons to Day 0 isolate). HPI = Hours Post Infection. F, H: Total virus production on Vero/TMPRSS2 cells measured until peak tire at 33°C (48 HPI peak) and 37°C (36 HPI peak) respectively, one-way ordinary ANOVA with Bonferroni’s multiple comparisons test (all comparisons to Day 0 isolate). I, K: Growth curves showing virus replication in hNECs at 33°C and 37°C respectively, 3 independent experiments, standard deviation shown on error bars, two-way repeated measures ANOVA with Bonferroni’s multiple comparisons test (all comparisons to Day 0 isolate). J, L: Total virus production on hNECs measured until peak tire at 33°C (120 HPI peak) and 37°C (72 HPI peak) respectively, one-way ordinary ANOVA with Bonferroni’s multiple comparisons test (all comparisons to Day 0 isolate).

There were replication differences in Vero/TMPRSS2 growth curves between Day 137 and the reference Day 0 isolate that were prominent at 33°C but less apparent at 37°C (Fig. 2 e, g). This temperature-dependent replication difference is highlighted by reduced total virus production of Day 137 isolate at 33°C but not 37°C (Fig. 2 f, h). Overall, Day 137 isolate shows the most attenuated phenotype compared to Day 0 isolate on Vero/TMPRSS2 cells, while isolates collected in the days before and after Day 137 virus show no attenuation.

### Primary respiratory epithelial cell related cultures have revealed differences between Alpha, Delta and

Omicron replication, but have not been previously used to investigate variants derived from immunocompromised individuals [49], [50]. The physiological relevance of polarized hNEC cultures can reveal virus fitness differences not apparent on widely used immortalized cell line models [26], [30], [51], [52], [53]. On hNECs, there were no differences in total virus production between the Patient 2 virus isolates at 33°C, though some timepoints displayed slight differences across the isolates (Fig. 2 i, j). However, the Day 137 isolate showed reduced total infectious virus production and a reduction in virus titres at multiple timepoints in hNEC cultures at 37°C (Fig. 2 k, l), with differences reaching an approximately ten-fold reduction in Day 137 isolate TCID_50_/ml versus Day 0 isolate at each timepoint between 24 to 48 hours post infection. Day 144 isolate showed reduced infectious virus production at early timepoints on hNECs at 37°C despite no significant attenuation at 33°C, or at either temperature on Vero/TMPRSS2 cells, suggesting that the few mutations that distinguish it from Day 134 isolate may affect the kinetics of infectious virus production, though not overall virus particle production (Fig. 1, Fig. 2k, l).

Overall, these results indicate temperature and cell culture-dependent differences in infectious virus production with Patient 2 isolates. Attenuation of the Day 137 isolate on hNECs at 37°C but not 33°C suggests that some later timepoint viruses in Patient 2 may have reduced fitness at temperatures corresponding to the lower respiratory tract.

### Patient 2 virus isolates do not show escape from neutralisation with convalescent plasma

Patient 2 received approximately weekly convalescent plasma from Day 103 onwards during their persistent infection [9]. Samples of plasma from Patient 2 or the convalescent plasma which Patient 2 initially received were no longer available and were not quantified for neutralising titre so we could not assess directly any escape from neutralizing antibodies in plasma collected over the course of the infection. As an alternative to assess escape from neutralizing antibodies, PRNTs were conducted using convalescent plasma from 8 donors across the US (with a known range of neutralising antibody titres), who were infected in the same time window as the Patients 2 and 3 to mimic the polyclonal antibody pressure present in the population during the period in which Patient 2 was shedding infectious virus [35]. Patient 2 Spike proteins do not contain RBD mutations, but N terminal domain (NTD) mutations (Fig. 3a) can increase resistance to neutralisation by vaccine-induced antibodies, as is the case with the Delta variant [12], [54]. There was no decrease in IC50 value for virus neutralization with any Patient 2 isolate (Fig. 3b), suggesting that escape from neutralisation by polyclonal antibodies was not a driving factor in the emergence of late timepoint viruses within Patient 2.

**Figure 3:**
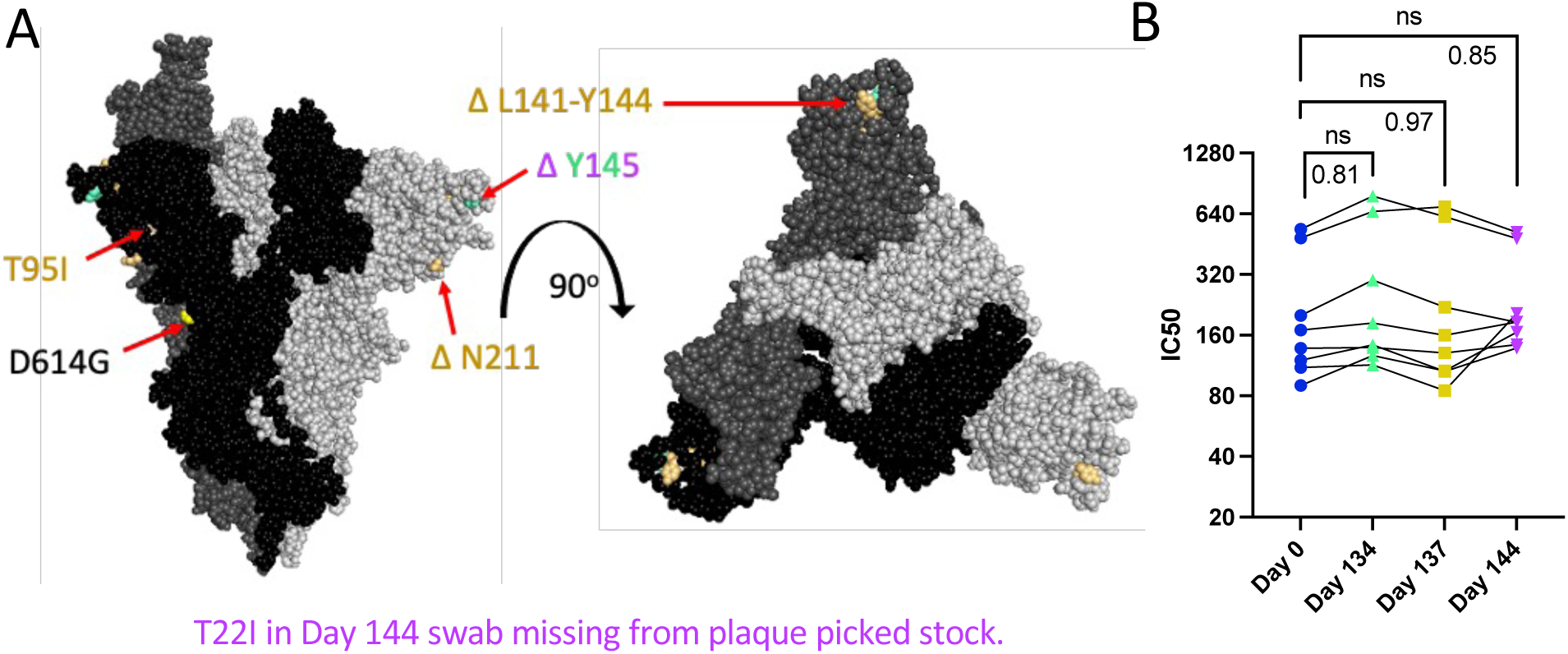
Spike mutations found on Patient 2 viruses do not correspond to major changes in neutralisation by convalescent plasma (B). Comparisons: * (Day 0 to Day 134), $ (Day 0 to Day 137), # (Day 0 to Day 144). “ns” (p > 0.05). A: Side and top view of the Spike trimer (each monomer in a different shade of grey, one RBD in up conformation), with Patient 2 virus-specific mutations displayed in corresponding colours, all virus Spikes contain D614G mutation (PyMOL, PDB: 7WZ2). B: PRNT IC50 values for Patient 2 viruses using 8 convalescent serum samples (each tested in duplicate) and graphed individually with lines connecting serum from the same individual, one-way repeated measures ANOVA with Bonferroni’s multiple comparisons test (all comparisons to Day 0 isolate).

### Patient 2 virus isolates do not show altered syncytia formation over the course of infection

All Spike mutations that appear in Patient 2 isolates are in the Spike protein’s NTD region (Fig. 3a). Mutations in the Delta variant Spike NTD increased cell-cell fusion, though the impact of NTD mutations on cell-cell fusion were dependent on Spike mutations outside of the NTD as well [54]. The Alpha variant’s H69/V70 deletion does not mediate immune escape, but increases cleaved Spike incorporation into the virus particle, resulting in an increased rate of syncytia formation [55]. To assess the impact of Patient 2 Spike mutations on Spike-induced cell-cell fusion, a two-colour syncytia assay was used (Fig. 4a) [56]. The level of Spike expression from the pCAGGS plasmid could drive differences in syncytia formation. To capture any differences in Spike expression from the pCAGGS-Spike plasmid preparations, flow cytometry to detect Spike at the cell surface of transfected Vero/TMPRSS2 cells was conducted in independent experiments (Supplementary Fig. 1). Flow cytometry indicated no statistical differences in percentage of Spike positive cells or mean fluorescence index (MFI) of Spike expression from the plasmids, (Supplementary Fig. 1b). For the syncytia assay, Vero/TMPRSS2/mCherry cells were transfected with pCAGGS plasmids containing Spike sequences identical to those in the corresponding Patient 2 isolates. Five hours after transfection, Vero/TMPRSS2/mCherry cells were mixed with CMFDA-treated TMPRSS2/ACE2 acceptor cells (green), and 24 hours after cell mixing, cells were fixed (Fig. 4a). Microscopy was then used to capture nuclei within the area of red/green overlap as an indication of fused red and green cells (Fig. 4b) [56]. There were no significant differences syncytia formation (Figure 4c), Spike surface expression or the percentage of Spike expressing cells (Supplemental Figure 1b) with Patient 2 isolates, indicating syncytia formation was not selected for in the evolution of Patient 2 viruses.

**Figure 4:**
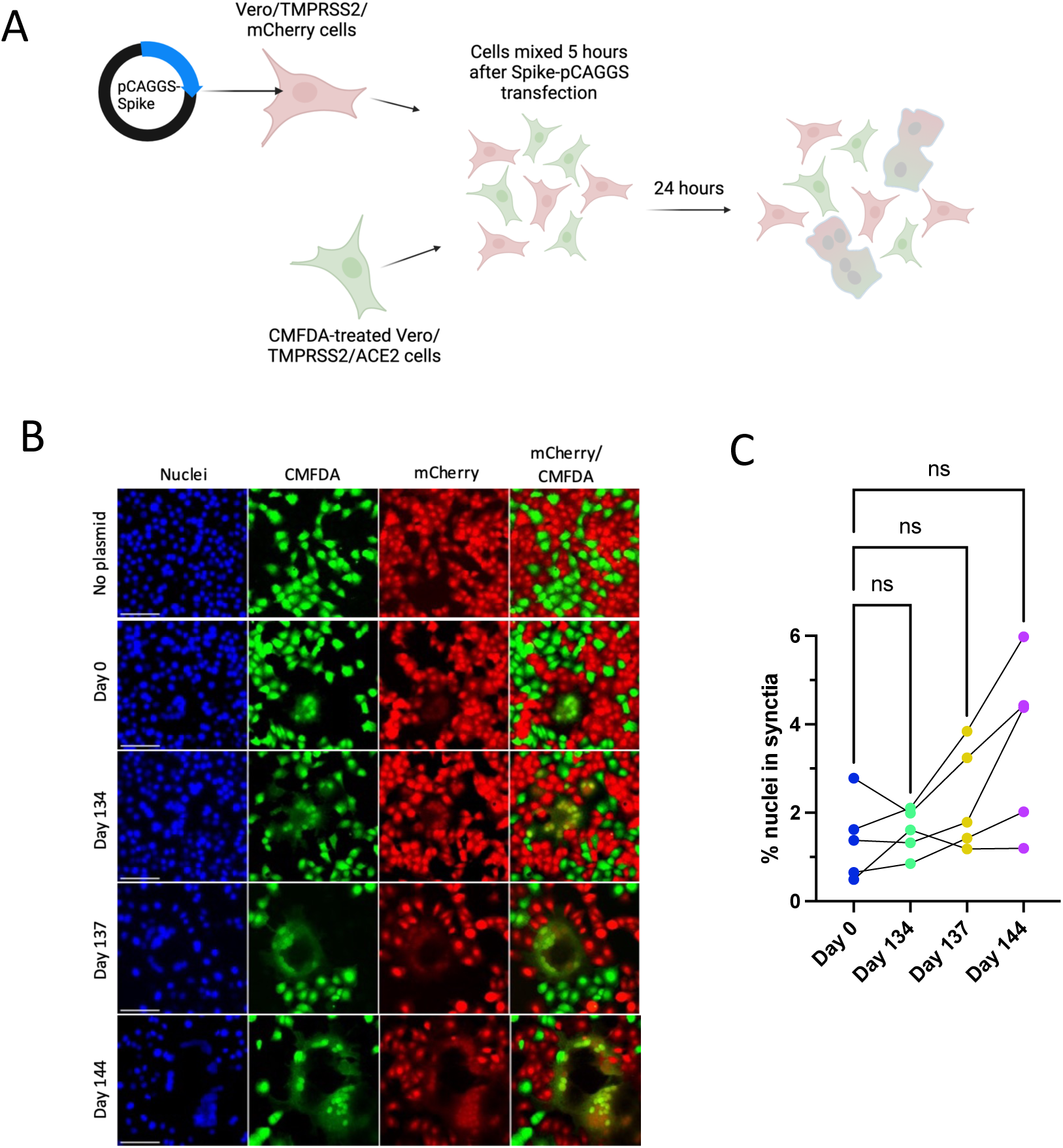
Patient 2 CHLA Spike mutations show trend towards increased syncytia formation versus Day 0 virus using two-colour syncytia assay. “ns” (p > 0.05). A: Overview of two-colour syncytia assay method (illustration created with BioRender). B: Example image of syncytia formation induced by each of the Patient 2 CHLA virus pCAGGS-Spike plasmids versus a no plasmid control. Scale bar = 100 μm. C: Percentage of nuclei within syncytia, 5 independent experiments graphed separately. One-way repeated measures ANOVA with Bonferroni’s multiple comparisons test (all comparisons to Day 0 isolate).

### Patient 3 virus isolates have different plaque sizes but no distinct replication differences on hNECs

At 33°C, Patient 3 Day 0 isolate had smaller plaques versus Day 139 isolate (Fig. 5 a, b). However, this size difference was reversed at 37°C, at which Day 139 plaques were visibly smaller (Fig. 5 c, d). Patient 3 Day 0 and Day 139 isolates show temperature-dependent differences in replication kinetics on Vero/TMPRSS2 cells, with Day 139 isolate reaching higher peak titers at 33°C and a faster peak titer at 37°C (despite smaller plaque sizes at 37°C) (Fig. 5 e,g). These differences in the kinetics of infectious virus production had no significant impact on total virus production (Fig. 5 f, h). However, virus replication differences were not apparent on hNECs at either temperature (Fig. 5i-l). Overall, these results suggest that differences in infectious virus production were not a major factor driving the selection for the combination of mutations found in Day 139 isolate in the persistently infected host, particularly when considering results from the more physiologically relevant hNEC model.

**Figure 5:**
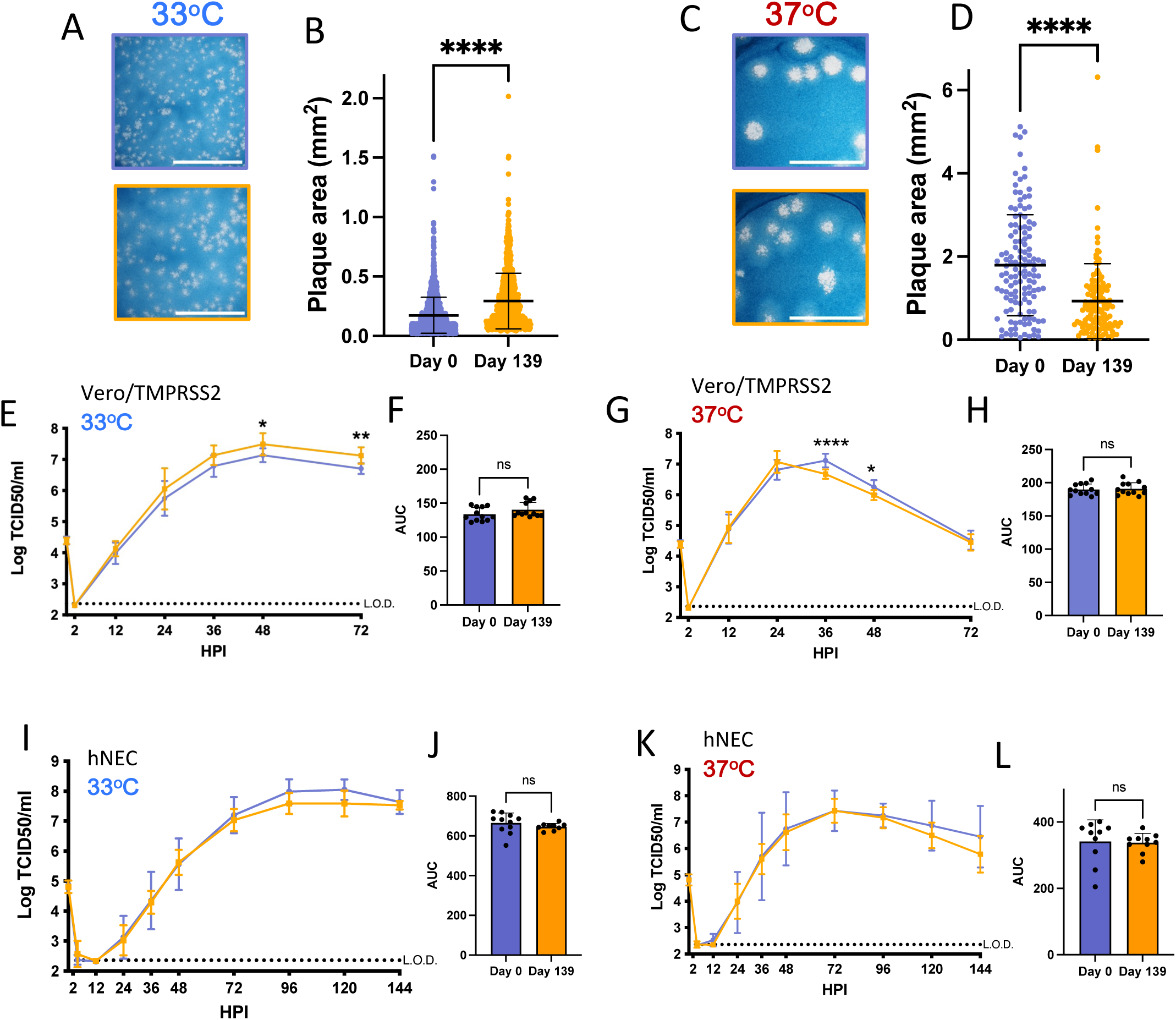
Virus isolated from Patient 3 later during the infection has distinct temperature-dependent plaque phenotypes but no significant replication differences. p values displayed as: “ns” p > 0.05,), * p < 0.05, ** p < 0.01, *** p < 0.001, ****p <0.0001. A, C: representative images of plaques in Vero/TMPRSS2 cells from each virus isolate at 33°C and 37°C respectively, scale bar = 10 mm. B, D: Quantified plaque sizes for > 889 (33°) and 129 (37°C) plaques per virus accumulated from 3 independent experiments on Vero/TMPRSS2 cells, unpaired t test. E, G: Growth curves showing virus replication on Vero/TMPRSS2 cells at 33°C and 37°C respectively, 3 independent experiments with four wells per virus per experiment, standard deviation shown on error bars, two-way repeated measures ANOVA with Bonferroni’s multiple comparisons test. F, H: : Total virus production on Vero/TMPRSS2 cells measured until peak tire at 33°C (48 HPI peak) and 37°C (36 HPI peak) respectively, unpaired t test. I, K: Growth curves on hNECs at 33°C and 37°C respectively, 3 independent experiments, standard deviation shown on error bars, two-way repeated measures ANOVA with Bonferroni’s multiple comparisons test. J, L: Total virus production on hNECs measured until peak tire at 33°C (120 HPI peak) and 37°C (72 HPI peak) respectively, unpaired t test.

### Patient 3 Day 139 virus has increased escape from neutralising convalescent plasma antibodies

Day 139 isolate mutations associated with escape from neutralising antibodies include ACE2 binding domain mutations V483A (23010 T/C) and E484Q (23013 G/C), and ΔL141-143 (21980 TTTTTGGTG/T) deletions known to abolish the binding of monoclonal neutralising antibody 4A8 (Fig. 6a) [12], [57], [58], [59]. Serum or plasma samples collected from Patient 3 during the time of infection were unavailable, and instead PRNTs were conducted using the same convalescent plasma panel as for Patient 2 isolates to assess neutralising antibody escape. In support of the cumulative effect of Day 139 Spike mutations on neutralising antibody evasion, there was an approximately 2.2 fold reduction in serum neutralizing activity across all 8 plasma tested against Day 139 isolate (Fig. 6b).

**Figure 6:**
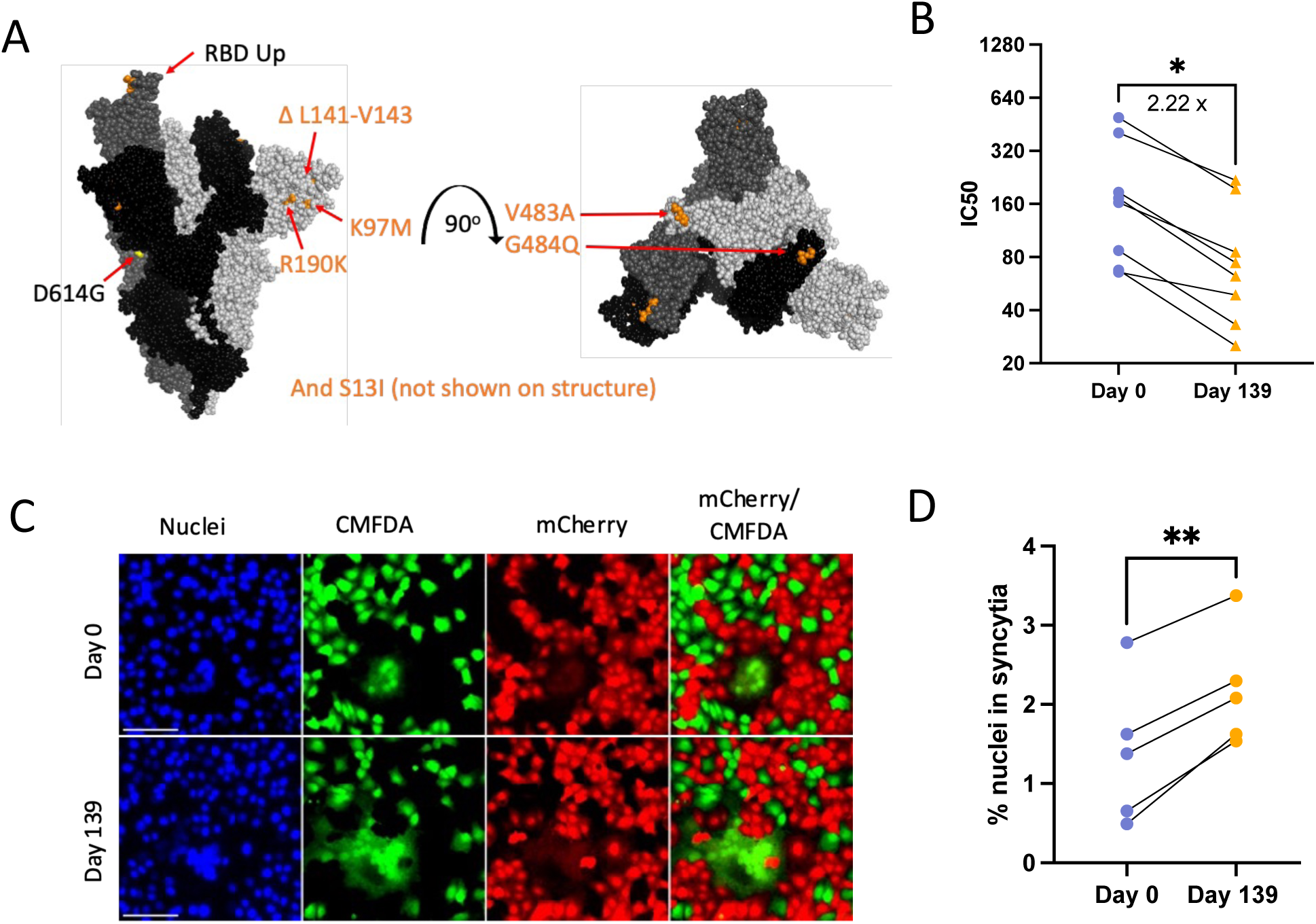
Patient 3 Day 139 virus shows significant escape from neutralising antibodies (B) and increased Spike-induced syncytia formation (D). p values displayed as: * p < 0.05, ** p < 0.01. A: Side and top view of the Spike trimer (each monomer in a different shade of grey, one RBD in up conformation), with Day 139 mutations displayed (no additional mutations apart from D614G on Day 0 Spike) (PyMOL, PDB: 7WZ2). B: PRNT IC50 values for Patient 3 viruses using 8 convalescent serum samples (each tested in duplicate) and graphed individually with lines connecting serum from the same individual. Fold change calculated from geometric means, paired 2-tailed t-test, * p < 0.05. C: Example image of syncytia formation induced by the Patient 3 CHLA virus pCAGGS-Spike plasmids versus a no plasmid control. Note Day 0 plasmid same as in Patient 2. Scale bar = 100 μm. D: Percentage of nuclei within syncytia, 5 independent experiments graphed separately. Statistics were performed on data pooled from all 5 experiments, paired 2-tailed t-test, ** p < 0.01.

### Day 139 Spike has increased syncytia formation versus Day 0 Spike

The Spike mutation E484K is known to increase Spike-ACE2 binding while reducing syncytia formation [60]. Deep mutational scanning maps indicate that E484Q also increases ACE2-binding affinity in the Wuhan-Hu-1 Spike background (though to a much lesser extent than E484K), and that V483A has no impact on ACE2-binding affinity [61]. However, the effects of these specific mutations on syncytia formation are unknown. Day 139 isolate Spike transfection consistently resulted in increased numbers of nuclei contained within syncytia versus Day 0 virus (Figs. 6 a, c, d). Consistent expression between Day 0 and Day 139 isolate Spike plasmids also means that expression differences were not a driving factor in differences in syncytia formation between these two Spike proteins (Supplementary figure 1b). Overall, results suggest that the unique combination of mutations found in Day 139 Spike drives increased syncytia formation along with escape from neutralising antibodies.

## Discussion

After approximately 140 days of a persistent SARS-CoV-2 infection, isolates from Patient 2 and Patient 3 were genotypically and phenotypically distinct, highlighting different potential trajectories for virus selection between ICPs. Mutations common to later timepoint Patient 2 and Patient 3 isolates include ΔL141-144 deletions in the Spike NTD, whereas Omicron-mirroring RBD mutations at V483 and E484 only appear in Patient 3 and not Patient 2 (Figs. 3a, 6a). In another example, the Envelope T30I (26333 C/T) mutation which has been observed in other case studies of persistently infected ICPs, features in Patient 3 Day 139 isolate but not in Patient 2 isolates [40], [62].

Differential plaque sizes between viruses can be driven by factors including virus replication rate, evasion of host antiviral responses, and induction of cell lysis on Vero/TMPRSS2 cells [63], [64]. Patient 2 isolates from all later timepoints have smaller plaque sizes versus Day 0 at 37°C, and Day 137 and Day 144 isolates also have smaller plaque sizes at 33°C. However, virus growth curves on Vero/TMPRSS2 cells only indicate significant attenuation of Day 137 isolate replication at 33°C, and Day 137 and Day 144 isolate attenuation on hNECs at 37°C. In contrast, differences in Patient 3 Day 139 isolate replication fitness were visible only on Vero/TMPRSS2 cells and not hNECs, and plaque size trends were inverted by temperature (Fig. 5). The trends for reduced replication fitness in Patient 2 isolates across plaque assays and growth curves suggest that persistent infections in ICPs do not necessarily select for variants that outcompete others in the respiratory tract on the basis of improved replication kinetics. The loss of replication fitness in Patient 2 isolates could also suggest that mutations that allow later timepoint viruses to persist over others in the respiratory tract could be detrimental to replication fitness *in vitro*. While replication fitness trends in Patient 3 isolates appear more nuanced with temperature-inverted trends in plaque-sizes, the lack of replication differences on hNECs at either 33°C or 37°C suggests that intra-host variant selection is not necessarily driven by variants outcompeting others at the level of replication. In addition, while our results indicate that mutations appearing in late timepoint viruses may not improve replication fitness *in vitro*, whether the variants appearing in Patients 2 and 3 show altered transmissibility is unknown.

The selection of antigenically distinct epitopes over the course of persistent infections in ICPs has been observed in SARS-CoV-2 and other RNA virus infections including norovirus [12], [26], [65]. Patient 2 isolates do not show increased escape from neutralisation by donor convalescent serum, with Day 134 isolate showing slightly increased susceptibility to neutralisation versus Day 0 isolate. While this patient received convalescent plasma, plasma IgG levels of SARS-CoV-2 specific antibodies were low and perhaps did not reach a concentration high enough to induce a selective pressure [9]. However, late time point isolate in Patient 3 (Day 139 isolate) had significantly increased escape from neutralising antibodies versus Day 0 isolate (Fig. 6). Patient 3 produced some SARS-CoV-2 specific antibodies, but these were dominantly IgM antibodies [9]. While IgM can be lower in affinity than IgG, its neutralising activity can often be stronger and broader than IgG with the exact same variable regions, which may explain the selective pressure which led to the emergence of neutralising antibody resistant virus in Patient 3 [27], [66], [67], [68]. Low neutralising antibody levels in other ICPs have likewise resulted in no antibody escape mutations appearing in other ICPs infected for 30 and 192 days, supporting the theory that selective pressure needs to reach a certain threshold before escape mutations are selected for [20]. This requirement for selection pressure is also reflected at the level of spread within a population, with the first SARS-CoV-2 immune escape variants Beta and Gamma appearing at significant levels only in late 2020 as the reinfection of immune populations became an increasing limitation to transmissibility [69].

A neutralizing antibody presence in Patient 3 could have driven the selection for escape variants, as reflected in both the appearance of V483A and E484Q mutations (two mutations common to persistent infections across ICPs) and an escape from neutralising antibodies in our serum panel [40], [62]. Critically, the Day 139 isolate shows escape from neutralising antibodies found in the serum of individuals acutely infected within the same timeframe as Patients 2 and 3, indicating that virus present within Patient 3 at day 139 may have had the ability to escape population immunity and cause reinfections if it had transmitted from Patient 3 [69]. Our results emphasise that over the course of persistent infections, some ICPs may be more likely to become a source of immune escape variants than others, and further work to characterise variant viruses in ICPs for immune escape is warranted. In addition, the RBD site mutations contained within Patient 3 Day 139 isolate may drive the difference in antibody escape trends between the Patient 2 and Patient 3 late time point isolates [59].

Cell-cell transmission within the respiratory tract may provide another means of antibody response evasion, as SARS-CoV-2 virions can infect neighbouring cells without becoming exposed to extracellular antibodies [70]. Patient 3 Day 139 isolate Spike consistently induced increased syncytia formation versus Day 0 Spike. Within the ICP host, Day 139 isolate mutations could have conferred increased immune escape through both neutralising antibody escape and increased cell-cell spread, building a larger landscape for variant selection in Patient 3 within which immune evasion but not virus replication differences could have been a key driver for virus selection [70]. Similarly, the Alpha variant of concern showed increased Spike-induced syncytia formation and transmission but limited immune escape, and had a concentration of NTD mutations (alongside furin cleavage site mutations) independently of immune escape mutations at the RBD [60], [71].

Temperature can affect virus replication kinetics and host cell responses to virus infections, and the temperature at which respiratory viruses replicate best can define their transmissibility. We and others have previously shown that physiological temperature ranges can alter influenza A virus and live attenuated influenza A virus replication kinetics on immortalized and primary cell cultures, and influenza B virus hemagglutinin protein expression is increased at cooler temperatures corresponding to the upper airway [51], [52], [53], [72], [73]. SARS-CoV-2 replication on hNECs is also variable at 33°C versus 37°C, and early A-lineage SARS-CoV-2 isolates show temperature-dependent differences in replication on both Vero/TMPRSS2 cells and hNECs [30], [74]. Comparisons using pseudoviruses bearing the SARS-CoV-2 Spike protein versus other human coronavirus Spikes using indicate that Spike is a key driver of coronavirus temperature preferences, and psuedovirus infectivity was heightened at 33°C in the case of SARS-CoV-2 but not SARS-CoV Spike [47]. The SARS-CoV-2 Spike D614G mutation also impacts SARS-CoV-2 infectivity at 33°C versus 37°C, underlining the potential for Spike protein mutations arising in ICPs to alter temperature preferences [47]. The Omicron variant showed increased airborne transmission versus earlier variants, improved replication in *in vitro* cell cultures mimicking the upper but not lower airways, and improved replication at 34°C versus 37°C unlike ancestral and Delta variants [75], [76], [77]. To our knowledge, this is the first characterization ICP-origin variant replication across physiological ranges of temperature. Understanding how SARS-CoV-2 variants may undergo continued temperature adaptation within a single ICP warrants further study, particularly as the temperature adaptation of viruses including SARS-CoV-2 and influenza can contribute towards changes in disease potential and possibly transmission efficiency.

The study of virus populations from ICPs is limited by the method of collection of virus isolates. Nasal swab sampling likely introduces biases towards viruses in the upper respiratory tract that replicate successfully at 33°C, and the true scale of SARS-CoV-2 diversity within the respiratory tract of persistently infected ICPs may not be captured by a nasal swab if other populations of SARS-CoV-2 variants are present within the same patient deeper within the respiratory tract. The potential for co-existing genetically distinct SARS-CoV-2 populations within the respiratory tract of ICPs is supported by the existence of genetically distinct influenza A populations between distinct lung lobes in infected ferrets [78].

During acute SARS-CoV-2 infections, intra-host virus diversity is very limited [4], [5], [79]. However, over the course of persistent infections in ICPs, many rounds of replication result in increased likelihoods of mutations in combination with selection [4]. In support of a diverse pool of viruses co-existing during the course of persistent infections, later timepoint virus isolates from Patient 2 have distinct genotypes and replication phenotypes despite being isolated from nasal swabs collected over a span of 10 days (Figs 1-4). Truong *et al.* conducted a time-resolved evolutionary rate estimation which suggested that Patient 2 Day 144 virus did not evolve sequentially from Day 134 virus [9]. Given the differences between Day 134, Day 137 and Day 144 isolate genotypes and replication phenotypes, it is plausible that major changes in variant composition within Patient 2 were driven by a rapid rise and fall of competing viral variants from a pool of variants found within the respiratory tract. Day 137 virus was rapidly replaced by Day 144 virus in Patient 2, which itself has a combination of mutations that slightly attenuate virus replication on hNECs at 37°C but not 33°C (Fig 2). Critically, Day 144 attenuation is minor compared to Day 137 isolate, suggesting that Day 144 isolate could also have outcompeted Day 137 virus in the respiratory tract environment. Swab samples from Patient 3 were less frequent, but given differences in variant composition between days 67 and days 68, it is likely that Patient 3 similarly harboured a diverse pool of virus variants within their respiratory tract over the course of a persistent infection [9].

The unique host environment in every ICP infection means that this group should be closely monitored for ongoing infectious virus shedding, particularly as little is known about how Omicron variants may change within a persistently infected host [25]. Surveillance and sequencing within this group has already foretold mutations later found within dominant global variants, and the characterisation of viruses isolated from two ICPs has demonstrated that these mutations can lead to differences in virus replication, syncytia formation and immune escape [40].

## Abbreviations

CM: Complete media
CPE: Cytopathic effect
hNEC: human Nasal Epithelial Cell
ICPs: immunocompromised patients
IM: Infection media
NTD: N-terminal Domain
PRNT: Plaque Reduction Neutralisation Test
RBD: Receptor Binding Domain
SNPs: single nucleotide polymorphisms

## Acknowledgement

This work was supported by National Institutes of Health (NIH) contracts N272201400007C and N75N03021C00045 for the Johns Hopkins Centers of Excellence in Influenza Research and Research, the U.S. Department of Defense’s (DOD) Joint Program Executive Office for Chemical, Biological, Radiological and Nuclear Defense (JPEO-CBRND), in collaboration with the Defense Health Agency (DHA) (contract number: W911QY2090012) with additional support from Bloomberg Philanthropies, State of Maryland and the Richard Eliasberg Family Foundation. The authors thank the patients who enrolled and participated in the project and are grateful for the efforts of the clinical coordination teams. We thank the laboratories of Sabra Klein, Kimberly Davis, Nicole Baumgarth, and Andrew Pekosz for discussion of data and future directions.

---

**Supplementary Figure 1:**
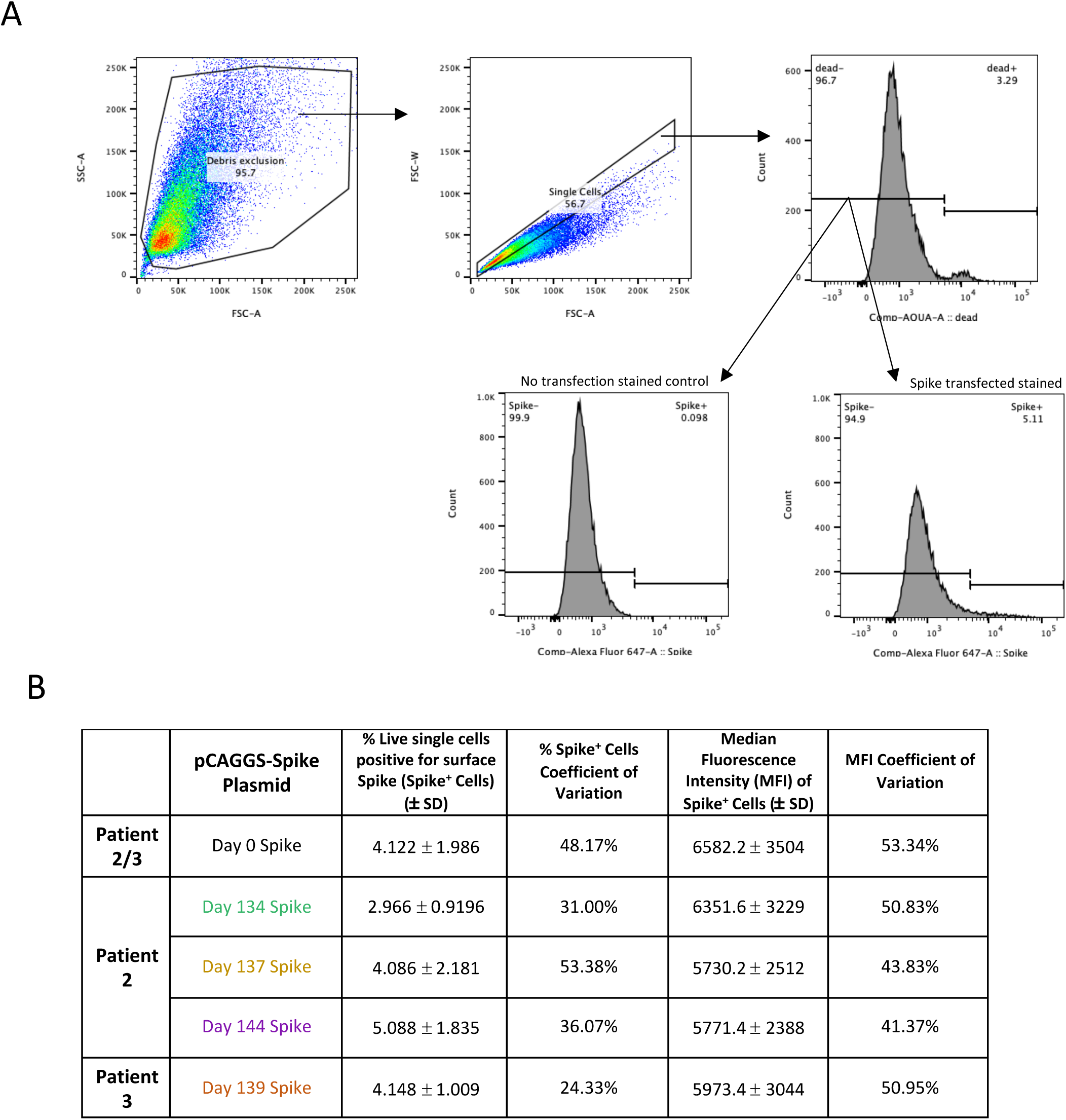
Flow cytometry gating strategy and results. A: Total cells were sequentially gated for debris exclusion, single cells and live cells (which do not take up the Live/Dead Fixable Aqua Dead Cell Stain), before gating for Spike positive cells. B: Mean percentage and Mean Fluorescence Intensity of cells positive for surface Spike for each pCAGGS-Spike plasmid, averaged from 5 independent flow cytometry experiments, ± standard deviation. One-way repeated measures ANOVA with Bonferroni’s multiple comparisons test (* p < 0.05) was performed on the data set and no significant differences were found between percent live Spike positive cells or MFI between Spike-pCAGGS plasmids.

**Supplementary table 1:**
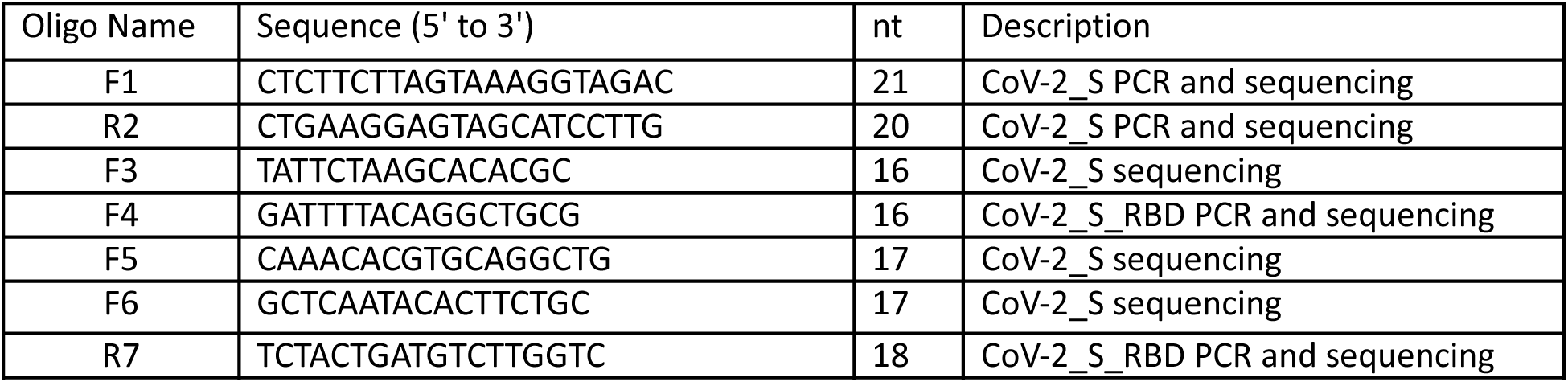
overlapping forward and reverse primers for Spike Sanger sequencing.

**Supplementary Table 2:**
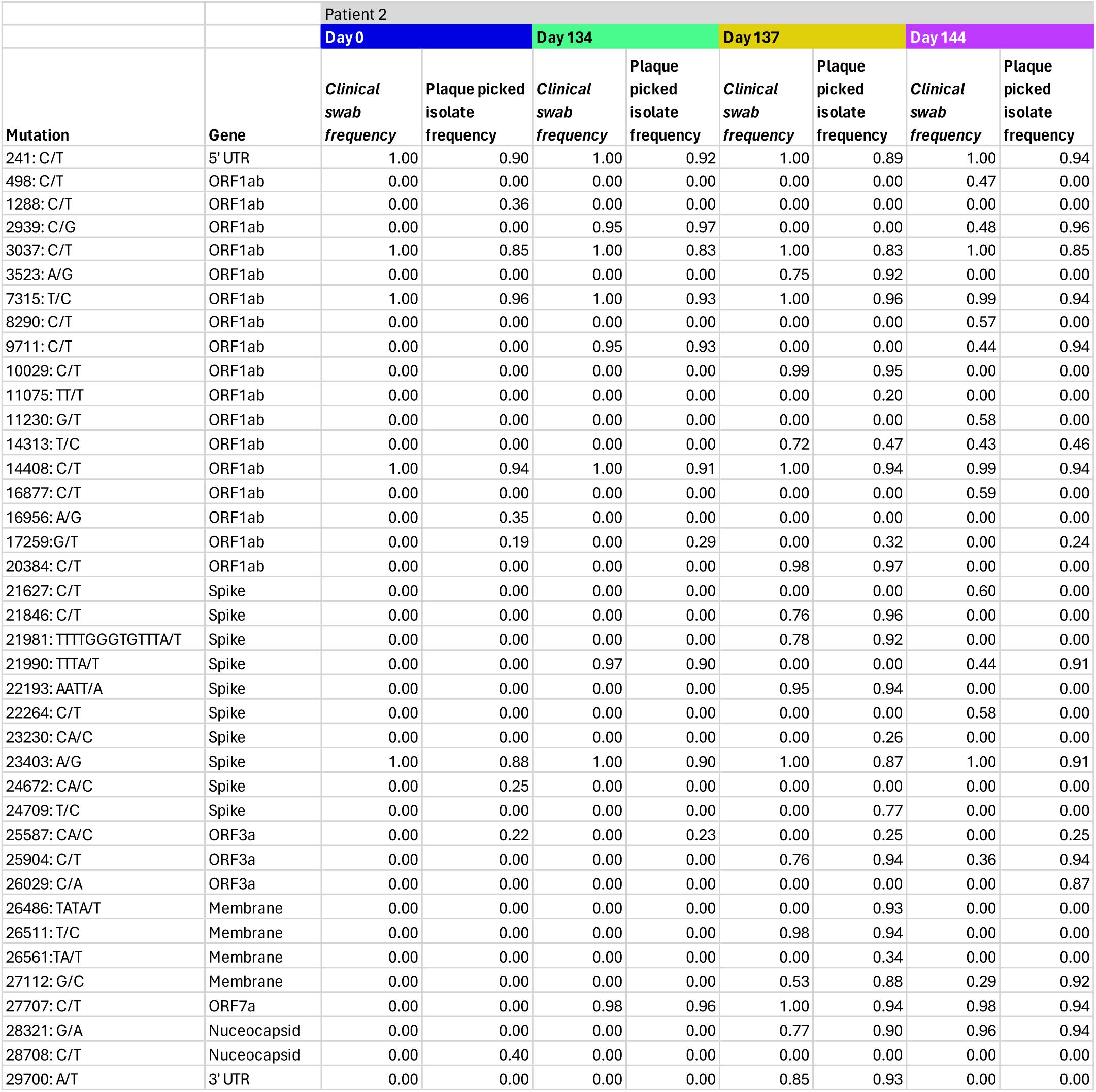
Comparison of mutation frequency for clinical swab samples versus plaque picked isolates for Patient 2 CHLA viruses, based on data from [[FINAL REFERENCE NUMBER FROM MANUSCRIPT]] and sequencing of plaque picked stocks.

**Supplementary Table 3:**
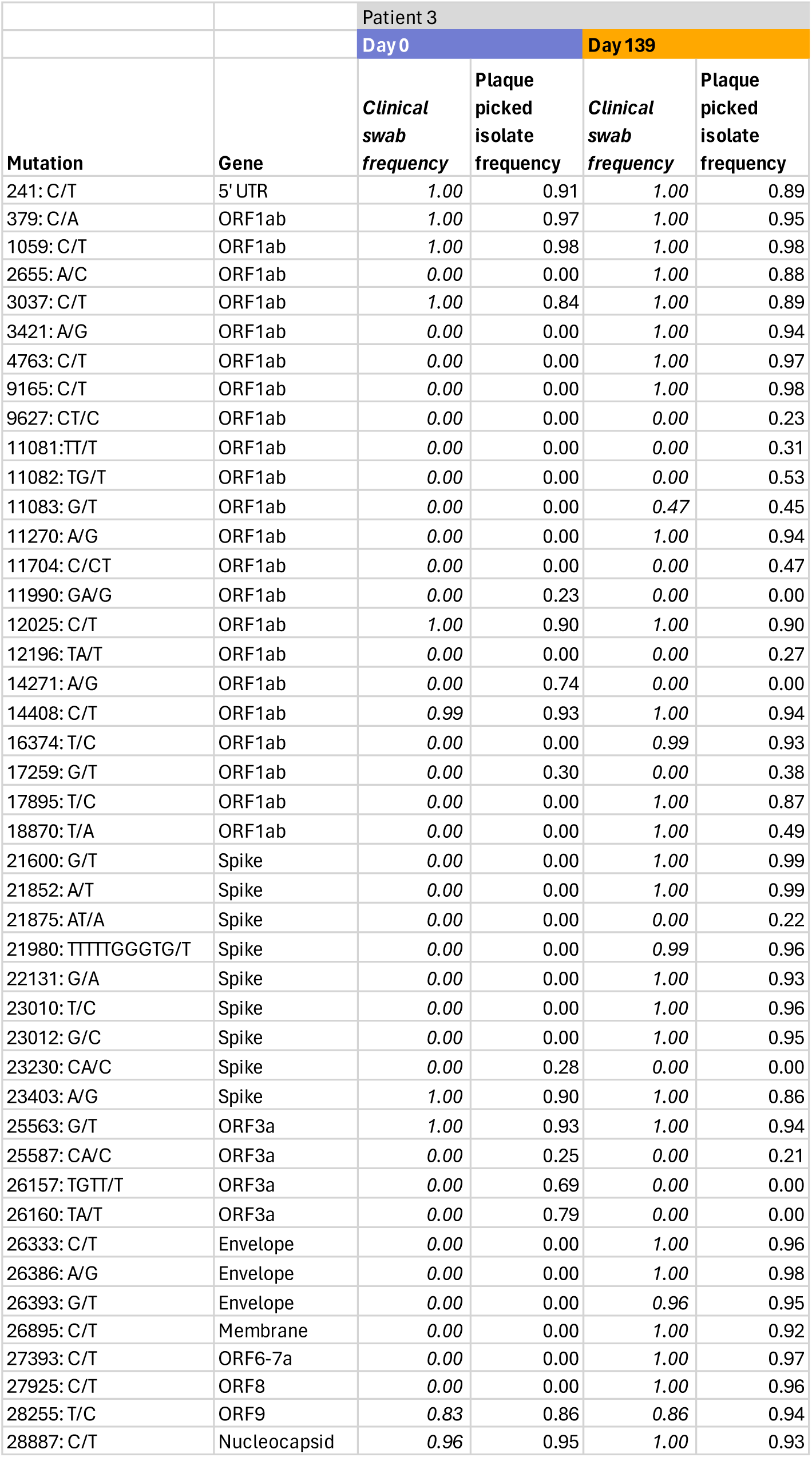
Comparison of mutation frequency for clinical swab samples versus plaque picked isolates for Patient 3 CHLA viruses, based on data from [[REFERENCE NUMBER FROM MANUSCRIPT]] and sequencing of plaque picked stocks.

## Notes

### Competing Interest Statement

The authors have declared no competing interest.

### Summary of Updates

several figures lost resolution or had missing panels in the initial submission. we have replaced the figure file with one that has all the correct panels in it, at good resolution.

